# FluidFM as a Tool to Study Adhesion Forces of Bacteria - Optimization of Parameters and Comparison to Conventional Bacterial Probe Scanning Force Spectroscopy

**DOI:** 10.1101/2019.12.19.882134

**Authors:** Linda Hofherr, Christine Müller-Renno, Christiane Ziegler

**Affiliations:** Department of Physics and Research Center OPTIMAS, University of Kaiserslautern, Erwin-Schrödinger-Strasse 56, 67663 Kaiserslautern, Germany

**Keywords:** FluidFM, SCFS, Immobilization, Bacterial adhesion

## Abstract

The FluidFM enables the immobilization of single cells on a hollow cantilever using relative underpressure. In this study, we systematically optimize versatile measurement parameters to improve the quality of force-distance curves recorded with a FluidFM. Using single bacterial cells, we show that Single Cell Force Spectroscopy experiments with the FluidFM lead to comparable results to a conventional Single Cell Force Spectroscopy approach using polydopamine for chemical fixation of a bacterial cell.

## Introduction

In this study, we focus on the FluidFM for assessing bacterial adhesion. Bacterial adhesion to surfaces is a key step in biofilm formation. Biofilms are applied in biofilm reactors for example in ethanol and butanol production or newly developed biofilm-based fuel cells [1–3]. Scanning Force Microscopy (SFM) and Scanning Force Spectroscopy (SFS) are versatile methods to study bacterial adhesion on surfaces. More specifically, single cell force spectroscopy (SCFS) is a prominent method to study the interaction of a single (bacterial) cell with a surface. Conventionally, a chemical functionalization of the SFM’s cantilever is used to immobilize a single bacterium for means of SCFS. However, due to different properties of bacteria, various functionalizations like polydopamine [4–6], poly-L-lysine [7–10], Cell-Tak [10,11] or glutaraldehyde [9,12,13] are used for different cells, making it necessary to test and find a suitable immobilization method for each cell type.

Some years ago, the FluidFM was introduced as an alternative method for the immobilization of cells on a cantilever [16]. The FluidFM is an add-on for conventional SFMs and allows the application of underpressure at the aperture of a hollow cantilever, which serves as micro- or nanopipette. For this purpose, the cantilever is filled with a liquid and coupled to a pumping unit, enabling a variation of the pressure inside the cantilever compared to the environment. By applying an underpressure relative to the environment, it is possible to draw a cell towards the opening of the cantilever and retain it there.

In this study, we determine how different measurement parameters like setpoint, z-speed, z-length, pause time, and relative underpressure influence the shape and quality of force-distance curves in a SCFS experiment using FluidFM. For this purpose, we studied two productive bacteria: The grampositive lactic acid bacterium *Lactococcus lactis*, which is applied in dairy industry as well as in the production of the antibacterial peptide nisin [18], and the gramnegative seawater bacterium *Paracoccus seriniphilus* which produces the enzyme serine dehydratase [19].

While focusing on the FluidFM, we implement measurement parameters for a SCFS measurement routine for *Lactococcus lactis*. A previous study showed that none of the typically used chemical immobilization methods leads to a reliable attachment of the cells without at the same time changing their surface properties [20]. Therefore, we used the FluidFM to physically immobilize the bacteria on the opening of a FluidFM cantilever and determined parameters for recording force-distance curves. Further, we used the well-studied gramnegative bacterium *Paracoccus seriniphilus* in terms of SCFS [17] to compare the immobilization by means of FluidFM with the well-established immobilization with polydopamine.

## Materials and Methods

### Bacterial Growth Conditions

The spherical, grampositive lactic acid bacterium *Lactococcus lactis* subsp*. lactis* was grown in DSMZ medium 11 (Leibniz Institute DSMZ – German Collection of Microorganisms and Cell Cultures). The spherical, gramnegative seawater bacterium *Paracoccus seriniphilus* was grown in complex medium as described elsewhere [21]. For both strains, precultures were incubated for 17 hours at 30° C. Afterwards, the precultures were diluted and kept at 30° C for another 24 hours before they were diluted in 0.9 % NaCl solution for subsequent SCFS measurements.

### Single Cell Force Spectroscopy

All measurements were performed with a Nanowizard III assembled with a FluidFM add-on (Cytosurge, Switzerland), positioned under an acoustic hood (both JPK Instruments AG, D) and mounted on an active vibration isolation system (Halcyonics-i4, Accurion GmbH, Germany) to minimize the effects of environmental vibrations. A droplet of diluted bacterial solution was placed on a microscope glass. For SCFS, a FluidFM nanopipette (Cytosurge, Switzerland) with an aperture of 300 nm and a nominal spring constant of 0.6 N/m was filled with glycerol, calibrated by the contact-based thermal noise method [14,15] and brought in close proximity to the glass surface. By applying a pressure of −800 mbar compared to the environment, a single bacterium was drawn to the aperture of the cantilever. For subsequent measurements, the negative pressure was reduced to smaller values.

Alternatively, All-In-One-Tipless Cantilevers A (BudgetSensors, Bulgaria) with an average spring constant of 0.2 N/m were cleaned and functionalized with polydopamine (for details, see [5]). After functionalization, the cantilever was pressed onto a single bacterium for 3 min with a setpoint of 2 nN to bind the cell to the cantilever.

All force-distance curves of *Paracoccus seriniphilus* were recorded with a setpoint of 2 nN, a z-length of 2 µm and a z-speed of 2 µm/s.

## Results and Discussion

### Influence of measurement parameters on the adhesion force of a blank nanopipette

We evaluated the influence of the setpoint, the z-speed and the applied pressure on the adhesion force of a blank FluidFM nanopipette on a glass surface. Figure 1 shows the course of the adhesion force as a function of the setpoint (a) and the z-speed (b) without pressure applied. Whereas the adhesion force increases with an increasing setpoint up to 20 nN, there is no clear trend for the z-speed.

**Figure 1:**
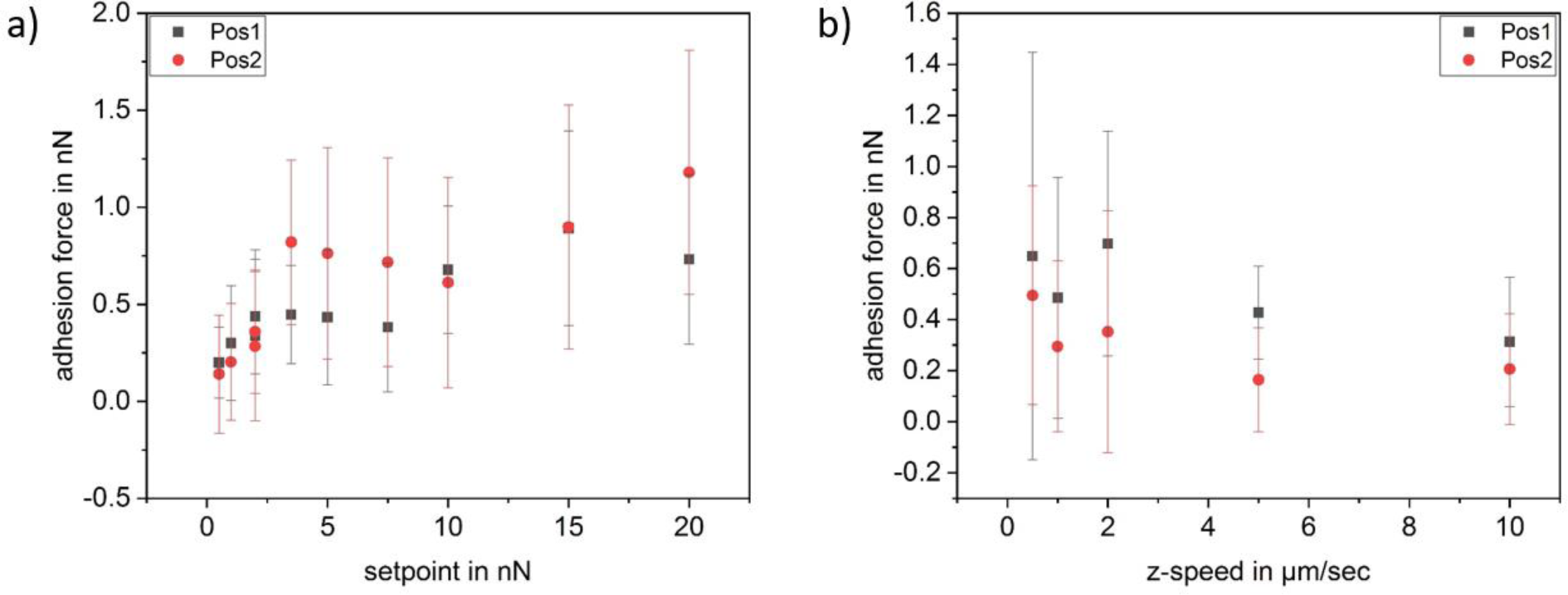
Adhesion force of a blank FluidFM nanopipette on glass, no pressure (0 mBar) applied; each data point comprises 25 force-distance curves a) as a function of the setpoint b) as a function of the z-speed

In comparison, fig. 2 shows the same measurements with an applied pressure of −50 mbar. Again, we see an increase of the adhesion force with the setpoint, but no clear trend regarding the z-speed. However, the absolute values of the adhesion force are slightly higher than without applied pressure.

**Figure 2:**
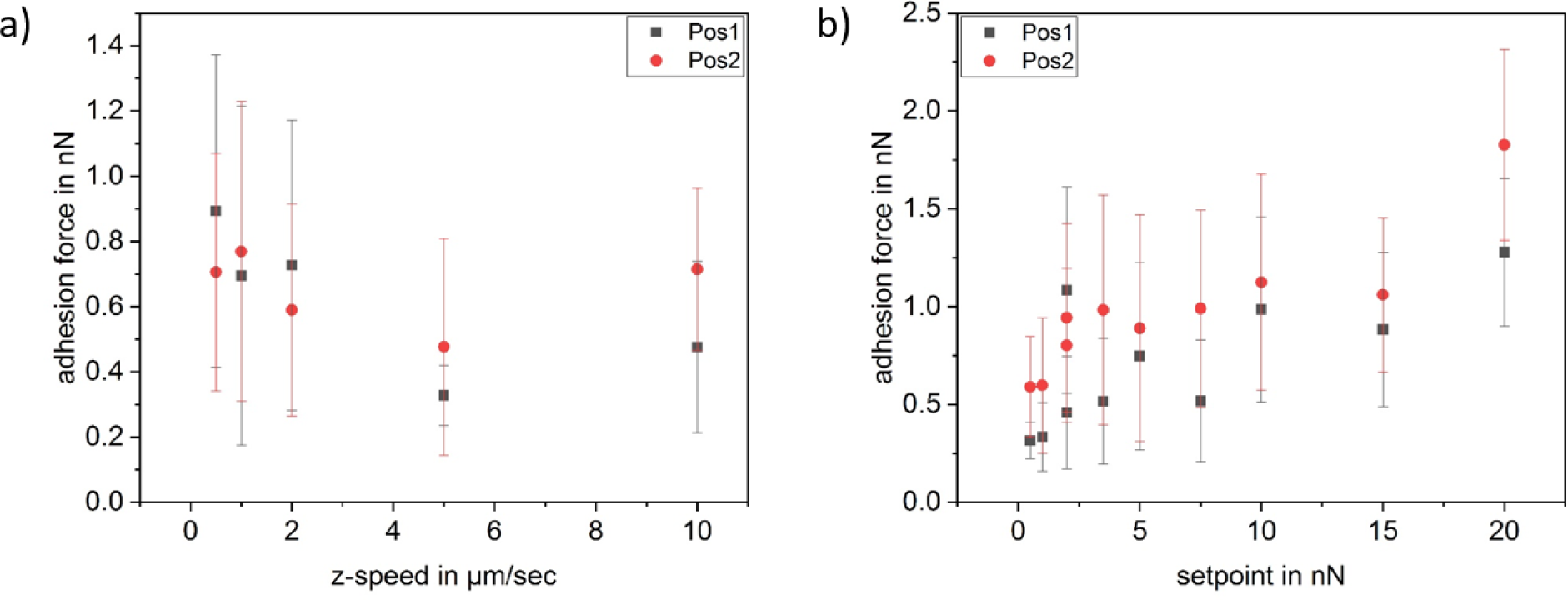
Adhesion force of a blank FluidFM nanopipette on glass, pressure of −50 mbar applied; each data point comprises 25 force-distance curves a) as a function of the setpoint b) as a function of the z-speed

Besides setpoint and z-speed, we determined the adhesion force of a blank nanopipette as a function of the applied pressure for two different sets of parameters (see fig. 3). The two sets of parameters match the ones used in the further course of the paper where the adhesion of single bacterial cells is investigated. In contrast to the subsequent measurements with bacteria, all recorded force-distance curves were of good quality for evaluation. Their shape is very similar to force-distance curves recorded with a conventional blank tip and shows a single sharp detachment event. Concerning the applied pressure, we see a clear trend of the adhesion force, namely a nearly linear drop with decreasing applied pressure up to 0 mbar. When the pressure rises above 0 mbar towards positive values, the adhesion force still decreases, but does not follow the linear course of the negative pressure values any more.

**Figure 3:**
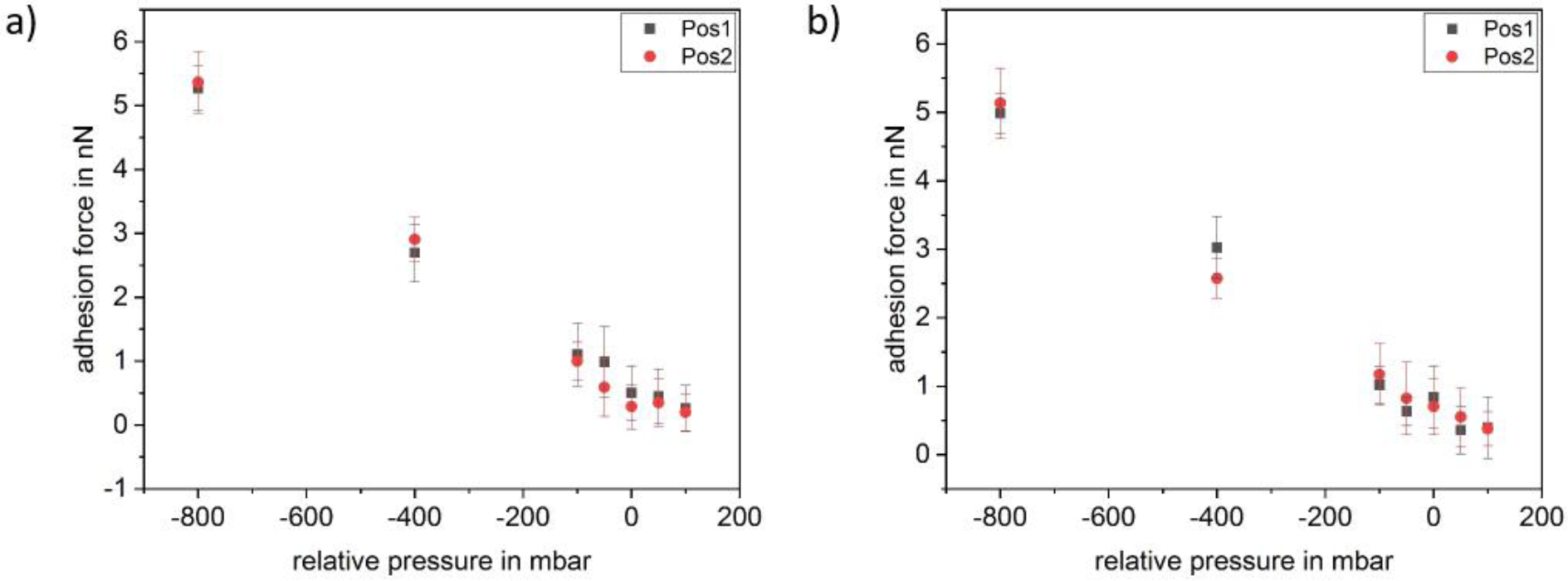
Adhesion force of a blank FluidFM nanopipette on glass as a function of applied pressure. Two different sets of parameters were used in a and b, each one representing the set of parameters for either *Paracoccus seriniphilus* (a) or *Lactococcus lactis* (b); each data point comprises 25 force-distance curves a) setpoint 2 nN, z-speed 2 µm/sec b) setpoint 5 nN, z-speed 5 µm/sec

### Influence and optimization of FluidFM measurement parameters

We investigated the influence of the parameters setpoint, z-length, z-speed, and pause time on the shape and quality of force-distance curves recorded by FluidFM. The investigation of each parameter was conducted using the formerly optimized parameters.

First, we studied the impact of the setpoint on the adhesion force and the quality of the force-distance curves. We found a clear correlation between the setpoint and the number of analyzable force-distance curves between 2 and 20 nN setpoint (2 nN, 3.5 nN, 5 nN, 7.5 nN, 10 nN, 15 nN, 20 nN; fig. 4). Further, we found the adhesion force increases with the setpoint, meaning the adhesion force strongly depends on the applied force.

**Figure 4:**
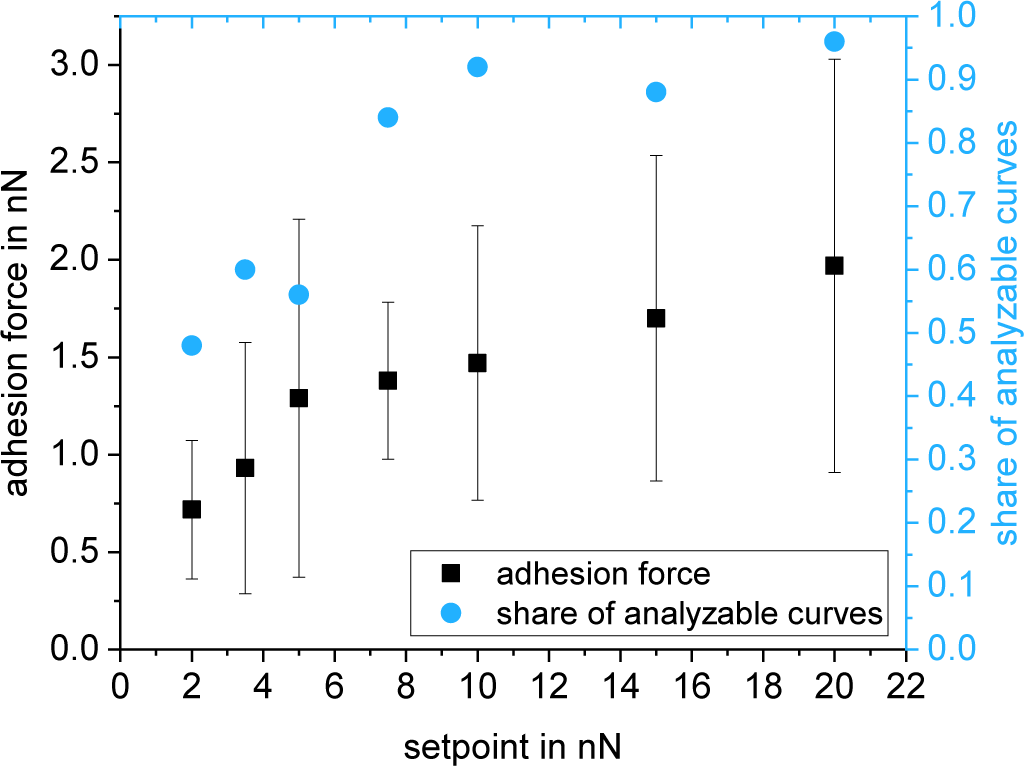
Adhesion force and share of analyzable force-distance curves of a single *Lactococcus lactis* cell on glass as a function of the setpoint; each data point comprises 25 force-distance curves

Fig. 5 shows an example of a force-distance curve recorded with a relatively small setpoint of 2 nN. Due to the hydrodynamic drift between approach and retract, the contact area of the retraction curve is not displayed completely for small setpoints. Further, the retract part shows a large drift in the area of the interaction-free sector, both being an indicator for a setpoint too small. However, the setpoint should be kept small in order to design the measurement as gentle as possible for the cells.

**Figure 5:**
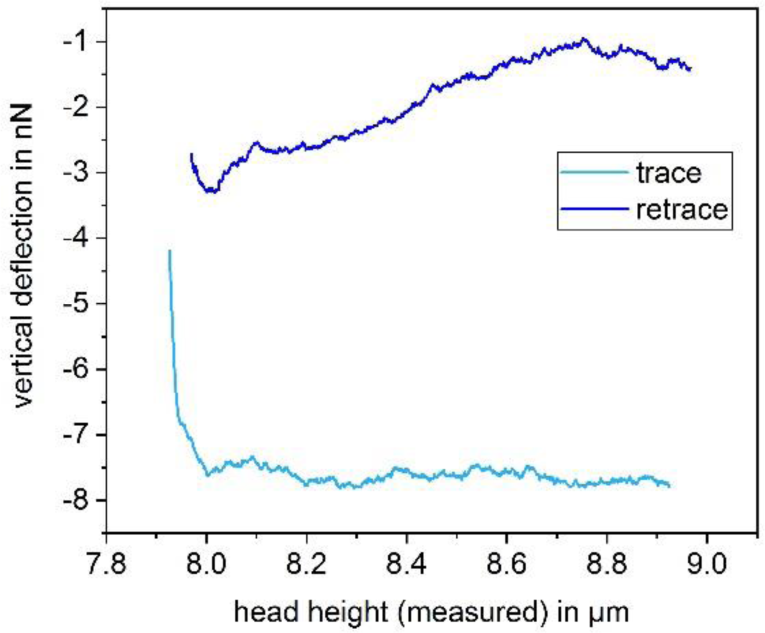
Force-distance curve of a single *Lactococcus lactis* cell on glass, recorded with the following parameters: setpoint 2 nN, z-length 1 µm, z-speed 2 µm/s, pause time 10 s

As for the z-length, the hydrodynamic drift increases with a growing z-length when comparing 0.5 µm, 1 µm and 2 µm z-length. Simultaneously, the stability of the baseline decreases. Figure 6 shows an example of a force-distance curve with an unstable baseline, recorded with a z-length of 1 µm. We determined that the quality of the force-distance curves increases with a decreasing z-length. Still, it is mandatory for the z-length to cover all rupture events. Especially when changing the surface material or the type of cells, it is mandatory to adjust the z-length to cover all rupture events.

**Figure 6:**
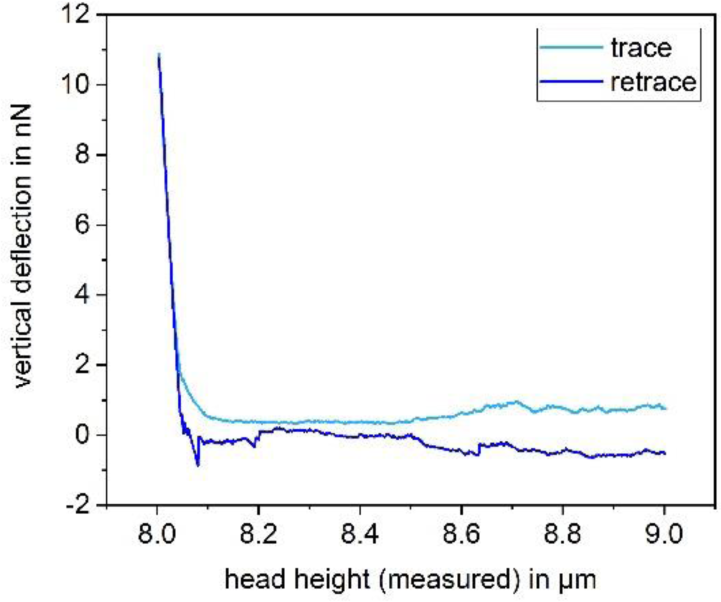
Force-distance curve of a single *Lactococcus lactis* cell on glass, recorded with the following parameters: setpoint 10 nN, z-length 1 µm, z-speed 2 µm/s, pause time 10 s

For the z-speed, we applied values between 0.5 and 10 µm/s (0.5 µm/s, 1 µm/s, 2 µm/s, 5 µm/s, 10 µm/s). Whereas small z-speeds cause the approach and retract curves to diverge (see fig. 7), this effect is smaller at higher z-speeds. Therefore, the z-speed should be increased to the point where this effect is minimized. For our setup, this was 5 µm/s. By further increasing the z-speed from 5 to 10 µm/s, we observed no further improvement of the force-distance curves.

**Figure 7:**
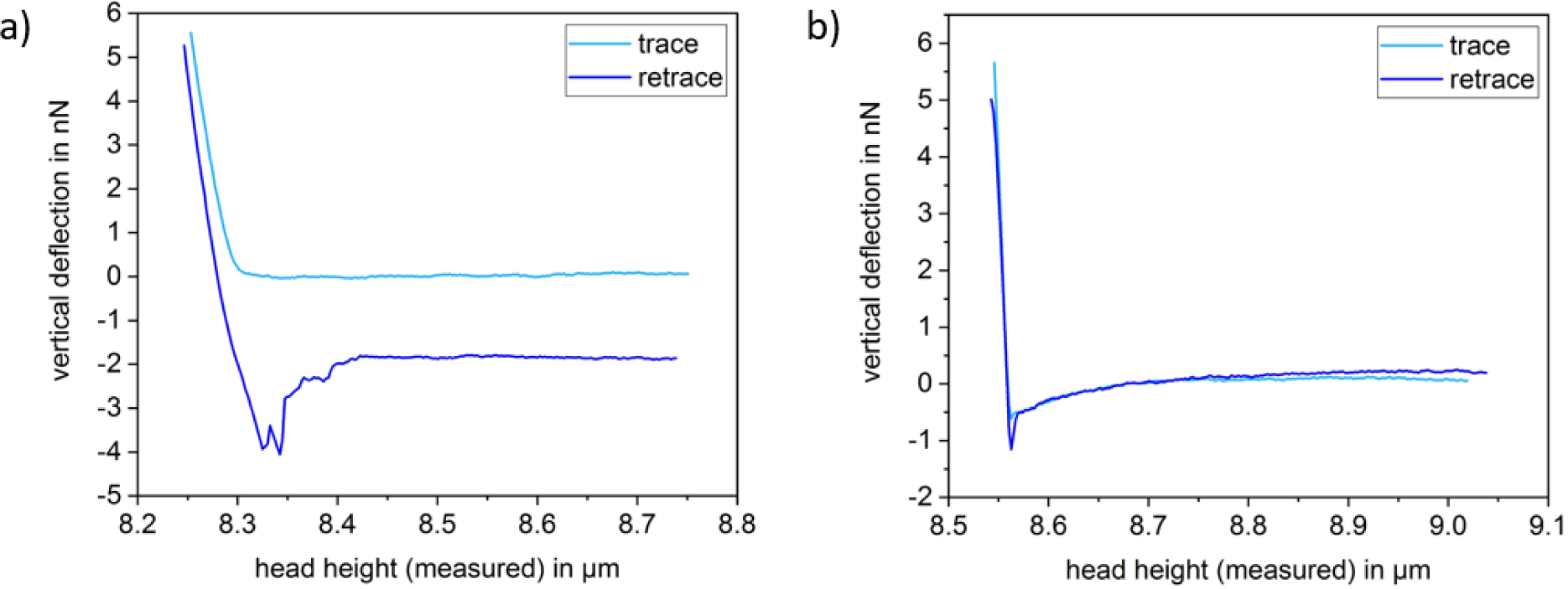
Force-distance curves of a single *Lactococcus lactis* cell on glass, recorded with the following parameters: a) setpoint 5 nN, z-length 0.5 µm, z-speed 5 µm/s, pause time 300 s b) setpoint 5 nN, z-length 0.5 µm, z-speed 5 µm/s, pause time 0.5 s time

Further, we studied the influence the pause time has on the shape of force-distance curves of *Lactococcus lactis*. For this purpose, we applied pause times between 0.5 and 300 s between approach and retract curves (0.5 s, 1 s, 2 s, 5 s, 10 s, 20 s, 60 s and 300 s). With a pause time larger than 2 s, typical bacterial rupture events are observable (see fig. 7a). For pause times shorter than 2 s, the curves change their appearance and show a single sharp rupture event and a curvature in the baseline (see fig. 7b). This behavior is reversible, meaning that by increasing the pause time, the bacterial rupture events reappear and the baseline straightens. Further, a longer pause time often correlates with an offset between approach and retract. However, this offset does not disturb the determination of the adhesion force as long as the curve is completely displayed.

The parameters with which the force-distance curves are recorded play an important role when determining adhesion forces by SCFS. By determining suitable parameters for recording force-distance curves with *Lactococcus lactis*, we found the adhesion force increasing with the setpoint (see fig. 4). Further, for *Lactococcus lactis* in combination with the FluidFM, we found a setpoint of 5 nN, a z-length of 0.5 µm and a z-speed of 5 µm/s to be appropriate parameters to determine the adhesion force on glass, thus showing that the FluidFM is suitable for the immobilization of a single *Lactococcus lactis* bacterium and subsequent SCFS measurements, which had not been possible in a reliable way without changing the cell’s properties previously.

For investigating the force-distance curves of the gramnegative bacterium *Paracoccus seriniphilus*, we applied a setpoint of 2 nN, a z-length of 2 µm and a z-speed of 2 µm/s since these parameters yield force-distance curves of good quality for both the FluidFM and the chemical immobilization approach using polydopamine (see next section). Concerning *Paracoccus seriniphilus*, we investigated how the applied pressure used for the immobilization by FluidFM influences the adhesion force between bacterium and surface. From fig. 8 we conclude that the influence of the applied pressure is small compared to a decrease of the adhesion force with a growing number of force-distance curves recorded. Referring to the literature, the decrease in the adhesion force for the repeated contact between cell and surface is due to irreversible bonds. Once broken, these bonds are unable to contribute to the adhesion force between cell and surface when recording subsequent force-distance curves, thus leading to a continuous decrease.

**Figure 8:**
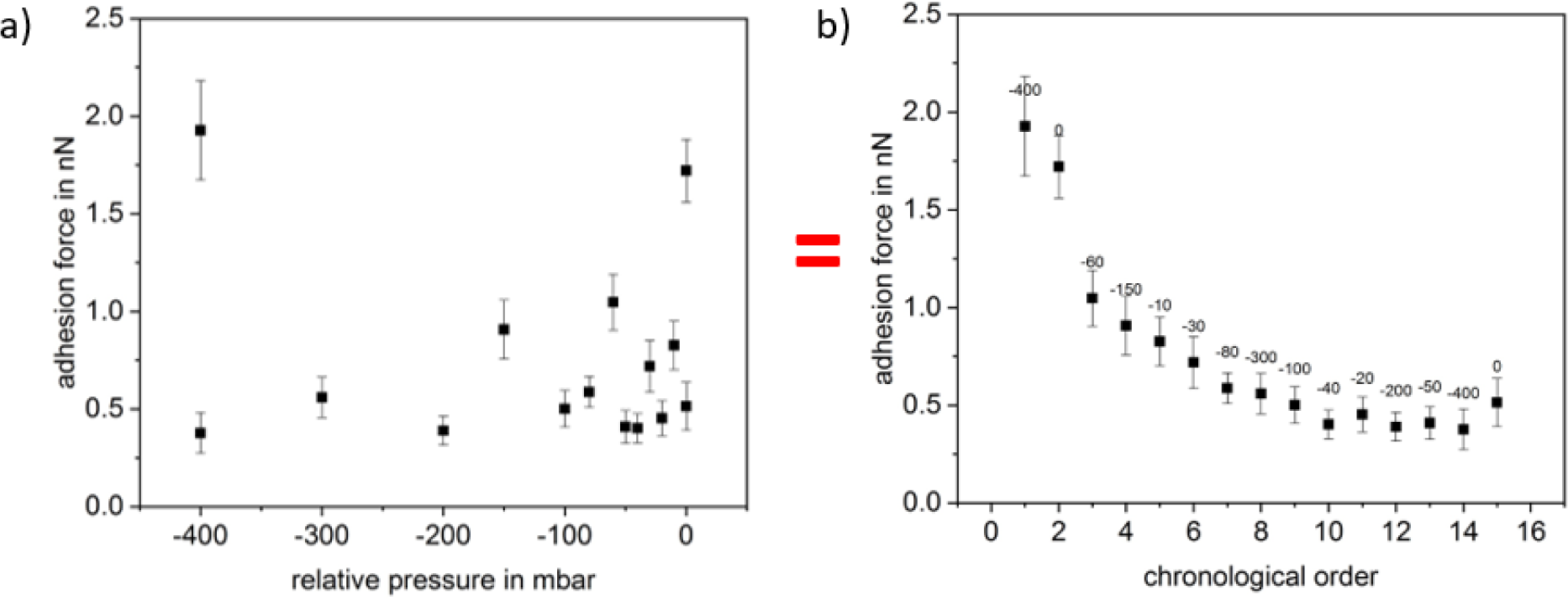
Adhesion force of *Paracoccus seriniphilus* on glass in a FluidFM experiment. Both, a and b, show the same data set, either sorted according to the applied pressure (a) or in chronological order (b); each data point comprises 25 force-distance curves

Once in fig. 8b the gradient due to the number of recorded force-distance curves decreased, the adhesion force for −20, −50 and 0 mbar exceeds the neighboring values which were recorded with higher suction force. Nevertheless, taking into account the error bars of the data points following −40 mbar, these changes could as well be due to the variance within the 25 force-distance curves recorded for each pressure. Since the cell is not held at the tip via underpressure when applying 0 mbar, we suggest that additional attractive forces act between the cells and the material of the nanopipette at the rim of the opening.

Further, since each data point in fig. 8 was recorded on a separate position on the glass surface, we conclude that the decreasing adhesion force due to repeated recording of force-distance curves at least partly results from changes of the interaction partner immobilized on the nanopipette. In fig. 8, this partner is the bacterial cell. To investigate if the changes in the cells are due to properties connected to the living nature of the bacterial cell, we expanded the measurements to polystyrene particles (Polybead Microspheres 1.00 µm, Polysciences, USA) instead of living cells. As can be seen in fig. 9, the decrease is not due to any malfunction or active adaptation of the biological component but can be observed for the elastic inanimate polystyrene particle as well.

**Figure 9:**
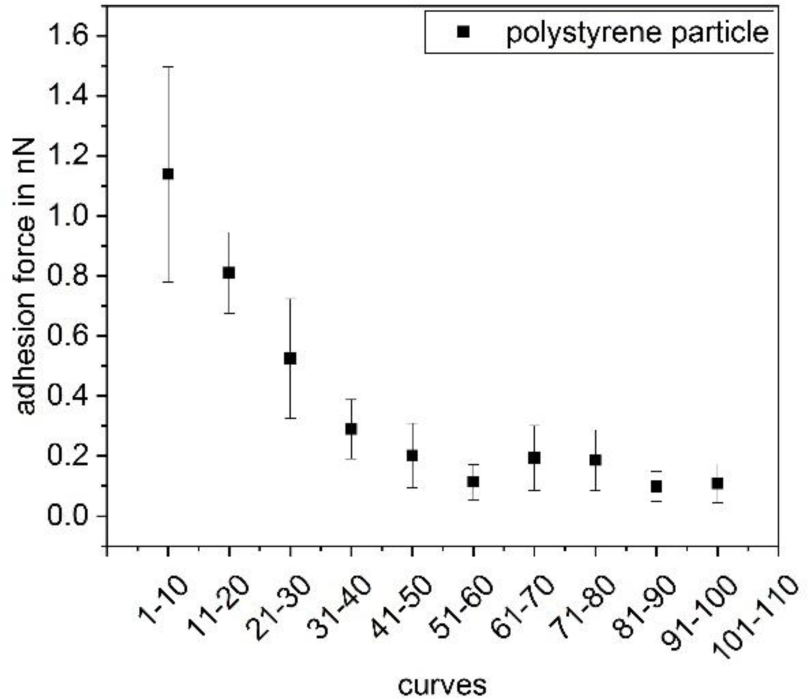
Adhesion force of a polystyrene particle as a function of the number of force-distance curves recorded previously

Besides studying the influence of the applied pressure, we recorded the adhesion force of a single *Paracoccus seriniphilus* cell as a function of the contact time between cell and a glass surface (see fig. 10). Whereas an increase of the adhesion force with the contact time was already shown elsewhere when *Paracoccus seriniphilus* is immobilized by polydopamine [17], we used the FluidFM for immobilization. To counteract the effect of decreasing adhesion forces with an increasing number of force-distance curves, we measured with increasing and subsequently decreasing contact times, thereby using the same cell. Especially in the beginning of the first measurement set, shown in black squares, the increase of the adhesion force due to an increasing contact time is strongly counteracted by the decrease due to the increasing number of recorded curves. This effect can be reduced by looking at the mean value of both measurement sets (triangles). However, the effect cannot be compensated completely.

**Figure 10:**
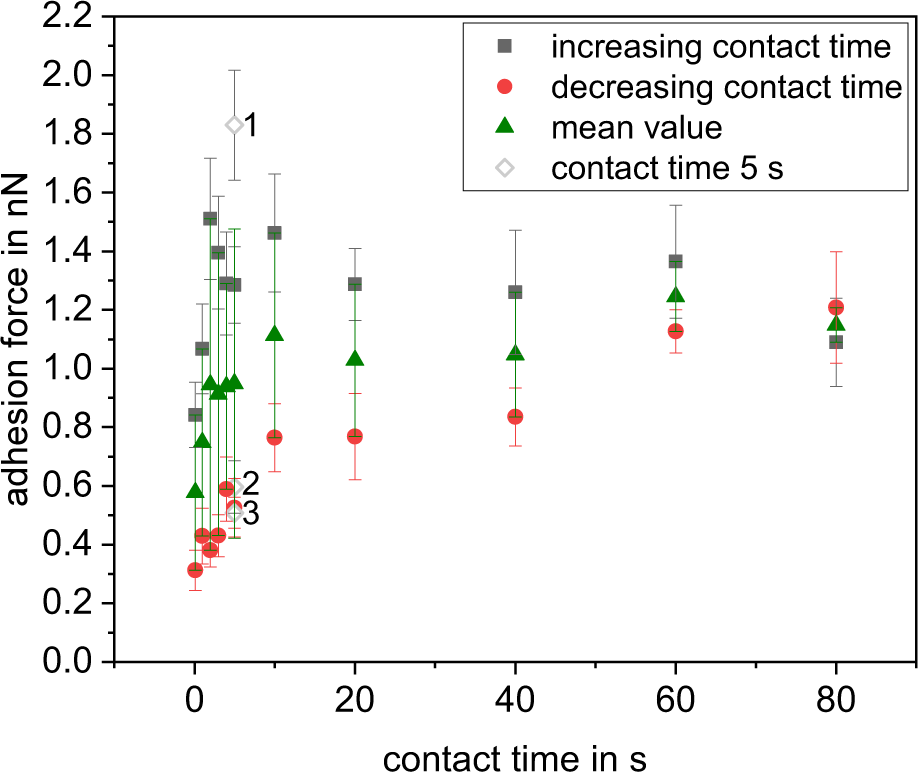
Influence of the contact time on the adhesion force of a single *Paracoccus seriniphilus* cell on glass. Each data point represents 15 force-distance curves. The data points depicted with void diamond at 5 s contact time were recorded at the beginning of the measurement (1), between the increasing and the decreasing contact times (2) and at the end of the measurement (3).

### Comparison between the immobilization of Paracoccus seriniphilus bacteria by polydopamine and FluidFM for SCFS

To assess if the immobilization with the FluidFM leads to different results compared to the conventional immobilization method using polydopamine, we investigated the adhesion force of *Paracoccus seriniphilus*, which is well-studied in terms of SCFS with polydopamine [17]. For this purpose, we compared the development of the adhesion forces of bacteria immobilized by the FluidFM with bacteria immobilized by means of the chemical functionalization of the cantilever with polydopamine.

Figure 11 shows the progression of the adhesion force of *Paracoccus seriniphilus* for multiple force-distance curves on glass, polished titanium and polished stainless steel, immobilized by polydopamine and FluidFM as well as a corresponding force-distance curve for both immobilization methods. 180 to 250 force-distance curves were recorded on one position on each surface, stainless steel being an exception in a way that the position was changed twice. For all cells and surfaces, the adhesion force of the bacterium decreases with a growing number of force-distance curves per position, regardless of the immobilization method. However, the extend of the decrease varies due to the individual cell that is used for the measurement. For example, the adhesion force decreases roughly linear on glass for both immobilization methods, as well as for one of the investigated cells on titanium if the cell is immobilized by FluidFM. On the other hand, the decrease on stainless steel is more pronounced within the first 100 force-distance curves. Further, since each data point is composed of ten force-distance curves, the adhesion force also decreases within the data points. The steeper decrease in the beginning of the measurement therefore causes higher error bars for the first data points compared to those of the data points recorded towards the end of the measurement.

**Figure 11:**
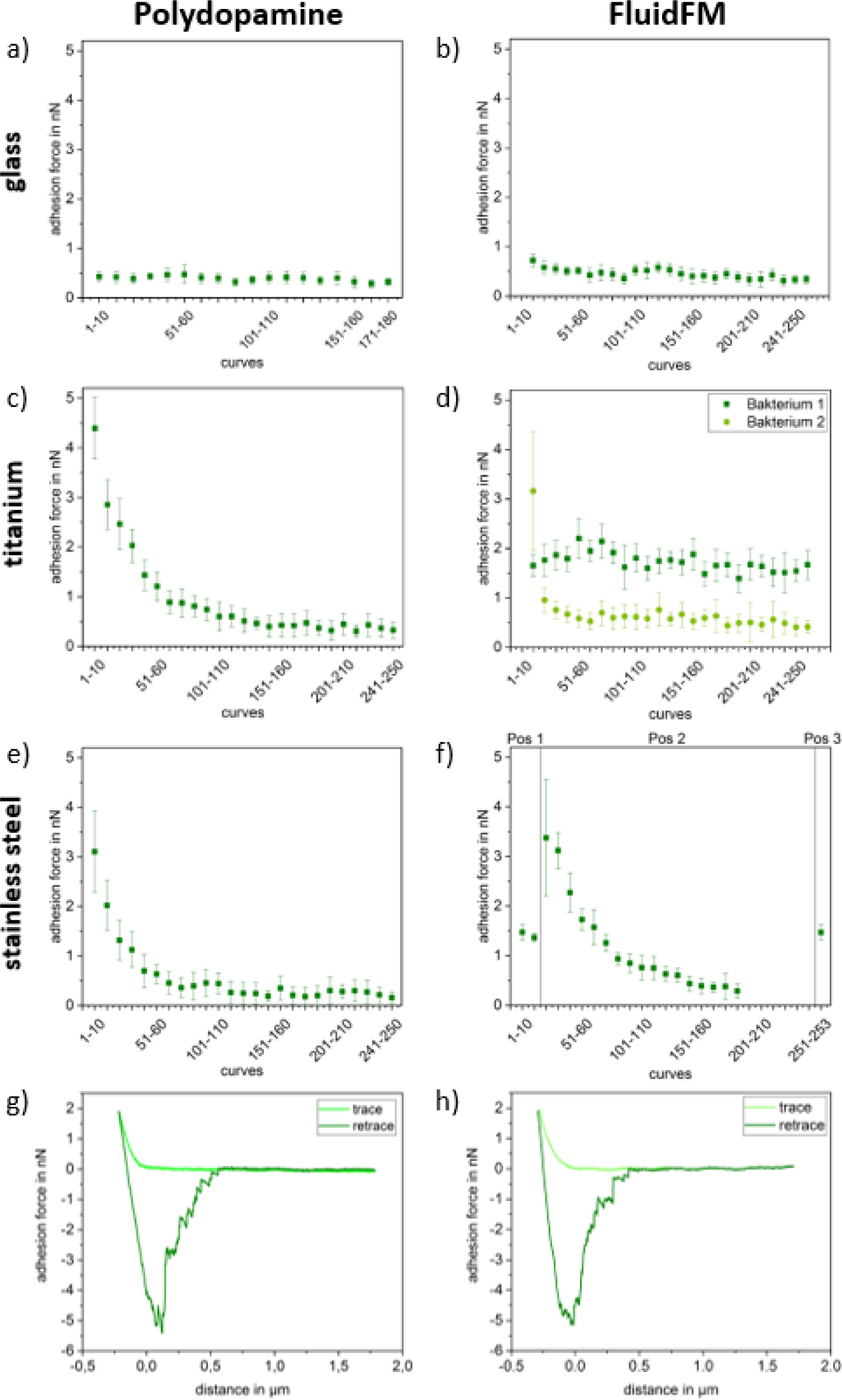
Adhesion forces (a-e) and force-distance curves (g, h) of *Paracoccus seriniphilus* on glass, titanium and stainless steel. The cell is immobilized on the cantilever either by polydopamine (left) or using the FluidFM approach (right). If not marked otherwise, all underlying force-distance curves of each plot were recorded on the same position. Measurement parameters were: setpoint 2 nN, z-length 2 µm, z-speed 2 µm/s

Further, taking into account the strong gradient between the data points in the beginning of the measurement set in fig. 9 and comparing them to the measurements on glass in fig. 11, this data supports the hypothesis that the different progression types observed in fig. 11 (linear and stronger in the beginning) are due to the use of different cells rather than to the immobilization method or the underlying substrate. Nevertheless, changes in the bacterial cells do not cause the changes solely. In fig. 11f, where the position was changed twice in the course of the measurement, a partial recovery of the original force value is observed, meaning that not only the bacterium but also the surface itself is lastingly influenced.

As for the comparison between the immobilization with polydopamine and FluidFM, the absolute force values are in the same range and it cannot be distinguished between both immobilization methods in this setup. Further, the force-distance curves look similar for both methods.

## Conclusions

We studied the influence of measurement parameters on FluidFM experiments and further showed that the results of a FluidFM experiment with a single bacterial cell lead to similar results compared to the conventional approach using polydopamine for immobilization. For a blank nanopipette, the adhesion force on a flat glass surface increases with the setpoint and when applying a suction force, whereas the z-speed shows no distinct influence.

We investigated the influence of the parameters setpoint, z-length, z-speed, pause time, and applied pressure on the shape and quality of force-distance curves of a single bacterial cell recorded by FluidFM. We found the adhesion force as well as the quality of the force-distance curves improve using higher setpoints in the range between 2 and 20 nN. Further, to obtain curves of good quality, a small z-length and a high z-speed should be applied. The applied underpressure had only a minor effect on the adhesion force of the cells. Even though the optimal parameters might differ using different bacterial strains, these general trends can help to find optimal parameters.

Further, we showed that the FluidFM is an appropriate and convenient method for the immobilization of *Lactococcus lactis*, a lactic acid bacterium that could not be immobilized reliably by conventional chemical functionalizations of the cantilever before. Further, by means of the well-studied *Paracoccus seriniphilus*, we showed that the physical immobilization using underpressure to hold the bacterium in place leads to the determination of adhesion forces similar to those obtained when applying polydopamine for immobilization. The reversible immobilization by the FluidFM allows for a quick change of cells and therefore enables to study a larger number of cells in less time compared to conventional chemical immobilization. Further, the FluidFM can be applied to immobilize various sorts of spherical bacteria, regardless of their outer membrane’s composition.

For the FluidFM as well as for polydopamine, we observed a continuous decrease of the adhesion force of a single cell with a growing number of force-distance curves, complicating the comparison between different environmental parameters in terms of adhesion forces and resulting from changes both of the cell and the underlying surface.

## Acknowledgement

This work is carried out in the Collaborative Research Center CRC 926 “Microscale Morphology of Component Surfaces”. Funding by Deutsche Forschungsgemeinschaft is gratefully acknowledged.

## References

[1] Zhang L, Zhou S, Zhuang L, Li W, Zhang J, Lu N, Deng L. Microbial fuel cell based on Klebsiella pneumoniae biofilm. Electrochem commun. 2008;10(10): 1641–1643.

[2] Nicolella C, van Loosdrecht MC, Heijnen SJ. Particle-based biofilm reactor technology. Trends Biotechnol. 2000;18(7): 312–320.

[3] Bühler K, Schmid A. Neue mikrobiologische Wege in der Chemie. BIOspektrum. 2011;17(5): 528.

[4] Kang S, Elimelech M. Bioinspired single bacterial cell force spectroscopy. Langmuir. 2009;25(17): 9656–9659.

[5] Beaussart A, El-Kirat-Chatel S, Sullan RMA, Alsteens D, Herman P, Derclaye S, Dufrêne YF. Quantifying the forces guiding microbial cell adhesion using single-cell force spectroscopy. Nat protoc. 2014;9(5): 1049.

[6] Beaussart A, El-Kirat-Chatel S, Herman P, Alsteens D, Mahillon J, Hols P, Dufrêne YF. Single-cell force spectroscopy of probiotic bacteria. Biophys J. 2013;104(9): 1886–1892.

[7] Bolshakova AV, Kiselyova OI, Filonov AS, Frolova OY, Lyubchenko YL, Yaminsky IV. Comparative studies of bacteria with an atomic force microscopy operating in different modes. Ultramicroscopy. 2001;86(1-2): 121–128.

[8] Schär-Zammaretti P, Ubbink J. The cell wall of lactic acid bacteria: surface constituents and macromolecular conformations. Biophys J. 2003;85(6): 4076–4092.

[9] Vadillo-Rodríguez V, Busscher HJ, Norde W, De Vries J, Dijkstra RJ, Stokroos I, Van Der Mei HC. Comparison of atomic force microscopy interaction forces between bacteria and silicon nitride substrata for three commonly used immobilization methods. Appl Environ Microbiol. 2004;70(9): 5441–5446.

[10] Meyer RL, Zhou X, Tang L, Arpanaei A, Kingshott P, Besenbacher F. Immobilisation of living bacteria for AFM imaging under physiological conditions. Ultramicroscopy. 2010;110(11): 1349–1357.

[11] Zeng G, Müller T, Meyer RL. Single-cell force spectroscopy of bacteria enabled by naturally derived proteins. Langmuir. 2014;30(14): 4019–4025.

[12] Razatos A, Ong YL, Sharma MM, Georgiou G. Molecular determinants of bacterial adhesion monitored by atomic force microscopy. PNAS. 1998;95(19): 11059–11064.

[13] Sheng X, Ting YP, Pehkonen SO. Force measurements of bacterial adhesion on metals using a cell probe atomic force microscope. J colloid interface sci. 2007;310(2): 661–669.

[14] Butt HJ, Jaschke M. Calculation of thermal noise in atomic force microscopy. Nanotechnology. 1995;6(1): 1.

[15] Hutter JL, Bechhoefer J. Calibration of atomic-force microscope tips. Rev Sci Instrum. 1993;64(7): 1868–1873.

[16] Meister A, Gabi M, Behr P, Studer P, Vörös J, Niedermann P., et al. FluidFM: combining atomic force microscopy and nanofluidics in a universal liquid delivery system for single cell applications and beyond. Nano letters, 2009;9(6): 2501–2507.

[17] Davoudi N, Huttenlochner K, Chodorski J, Schlegel C, Bohley M, Müller-Renno C, et al. Adhesion forces of the sea-water bacterium Paracoccus seriniphilus on titanium: Influence of microstructures and environmental conditions. Biointerphases, 2017;12(5): 05G606.

[18] Kuipers OP, Beerthuyzen MM, de Ruyter PG, Luesink EJ, de Vos W. Autoregulation of nisin biosynthesis in Lactococcus lactis by signal transduction. J Biol Chem. 1995;270(45): 27299–27304.

[19] Pukall R, Laroche M, Kroppenstedt, RM, Schumann P, Stackebrandt E, Ulber R. Paracoccus seriniphilus sp. nov., an L-serine-dehydratase-producing coccus isolated from the marine bryozoan Bugula plumosa. Int J Syst Evol Micr. 2003;53(2): 443–447.

[20] Hofherr L, Chodorski J, Müller-Renno C, Ulber R, Ziegler C. Comparison of Versatile Immobilization Methods for Gram-Positive Bacteria on a Silicon Cantilever. phys status solidi (a). 2018;215(15): 1700846.

[21] Davoudi N, Müller-Renno C, Ziegler C, Raid I, Seewig J, Schlegel C, et al. Nanomechanical properties of the sea-water bacterium Paracoccus seriniphilus—A scanning force microscopy approach. Biointerphases. 2015;10(1): 019004.

